# Phosphoproteomic Insights into TRPV4–AMPK Signaling Axis in the Choroid Plexus Epithelium: Implications for Therapeutic Targeting in Hydrocephalus

**DOI:** 10.64898/2026.07.13.738282

**Authors:** Gowthami Mahendran, Maryam Torabi, Bonnie Blazer-Yost

## Abstract

**Background:** Hydrocephalus is characterized by abnormal accumulation of cerebrospinal fluid (CSF) due to disrupted secretion, circulation, or reabsorption. CSF homeostasis is regulated by ion and water channels on choroid plexus epithelial (CPe) cells, including the mechanosensitive cation channel, transient receptor potential vanilloid 4 (TRPV4). Although TRPV4 antagonism prevents hydrocephalus progression in rats, how TRPV4 activity modulates CSF production remains unclear.

**Methods:** Because TRPV4 function is phosphorylation-dependent, we examined its activation (GSK1016790A) and inhibition (RN1734) and the downstream signaling alterations in human choroid plexus papilloma cells (HIBCPP) using phosphoproteomic mass spectrometry.

**Results:** Our phosphoproteomic analysis revealed significant changes in kinases and tight junction (TJ) proteins regulating epithelial barrier integrity. TRPV4 activation altered TJ proteins such as Zonula Occludens-1 (ZO-1), and Claudin-7 (CLDN7) and AMP-activated protein kinase (AMPK), phosphorylation, with a notable foldchange increase of AMPK inhibitory phospho form (Ser496). Further our electrophysiological and biochemical analyses demonstrated that AMPK activation followed by TRPV4 stimulation increased AMPK Ser496 phosphorylation, and epithelial permeability was subsequently elevated, although it was statistically not significant. In contrast, AMPK inhibition prior to TRPV4 activation resulted in the opposite effect with substantial epithelial barrier tightness which was evident from the ZO-1 expression. Moreover, a pharmacologically available anti-diabetic drug, metformin exhibited a similar trend like Compound C in reducing the barrier epithelial permeability after TRPV4 agonist treatment, highlighting that metformin promotes barrier tightness via an AMPK-dependent pathway coupled to TRPV4 signaling.

**Conclusions:** Overall, these findings identify an AMPK–TRPV4 signaling axis that modulates epithelial permeability and may influence CSF regulation, highlighting a potential therapeutic target for hydrocephalus.

## Introduction

The choroid plexus epithelium (CPe), present in the brain ventricles, is a specialized barrier epithelial monolayer surrounding a fenestrated capillary network and serves as the primary site of CSF production ^1^. CSF plays a critical role in maintaining intracranial pressure and central nervous system (CNS) homeostasis. The CPe forms the blood–CSF barrier (BCSFB) through tight junction (TJ) complexes between adjacent epithelial cells, and thereby tightly regulating transepithelial movement of ions and water to control CSF volume ^2–4^. Disruption of CPe function or barrier integrity can alter CSF dynamics and contribute to neurological disorders such as hydrocephalus, which is a multifaceted neurological condition characterized by the abnormal accumulation of CSF within the brain ventricles and increased intracranial pressure ^5,6^. This excessive fluid buildup leads to ventricular enlargement, disrupting delicate intracranial dynamics, impairing normal brain function and can give rise to a wide spectrum of neurological consequences, including cognitive and motor impairments as well as visual and gait disturbances ^7–10^. Emerging evidence from experimental models of hydrocephalus indicates that alterations in epithelial barrier integrity and ion transport at the CPe contribute to these pathological changes, underscoring the need to elucidate the molecular mechanisms governing these processes ^2,3,11–13^.

The CPe barrier function is maintained by highly specialized TJ complexes that form the structural basis of the BCSFB. These junctions are composed of transmembrane proteins, including claudins, occludins, and junctional adhesion molecules which assemble into continuous TJ strands that tightly regulate the paracellular movement of ions across the cells, defining a selective permeability of the epithelium ^14^. The cytoplasmic domains of these TJ proteins are anchored to the actin cytoskeleton through scaffolding proteins of the zonula occludens (ZO) family, particularly ZO-1, which organize junctional architecture and integrate signaling pathways ^15^. These CPe TJs are dynamic structures that can respond to intracellular signaling cascades, post translational modifications such as phosphorylation, and mechanical forces, regulating the epithelial permeability. Importantly, TJs also preserve epithelial polarity to ensure proper localization and function of ion transporters that drive CSF secretion. Disruption of TJ organization by inflammation, oxidative stress, infection or altered signaling can increase epithelial permeability, leading to abnormal ion fluxes, and dysregulated CSF production ^16^. However, the molecular sensors and signaling mechanisms controlling TJ remodeling in the CPe remain poorly understood.

One candidate mediator identified in the apical membrane of CPe that regulates the CSF production is the transient receptor potential vanilloid 4 (TRPV4) ^11,17^, which is a mechanosensitive, nonselective cation-permeable ion channel that is activated by osmotic stress and mechanical stimuli and, when activated, results in an increase in intracellular Ca^2+^ ^18^. As a mechanosensitive cation channel, TRPV4 has been shown to regulate cytoskeletal remodeling and intercellular junction organization in epithelial and endothelial cells ^19–21^. TRPV4 links mechanical or osmotic stimuli to intracellular calcium signaling to coordinate the movement of ions and water into and out of the cells to maintain the CSF dynamics ^22^. Studies show that targeting TRPV4 pharmacologically in the choroid plexus may attenuate CSF overproduction and ventricular enlargement in an experimental rat hydrocephalus model, highlighting its critical role in maintaining CSF homeostasis ^17^.

To study the role of CPe in CSF regulation, the HIBCPP cell line, derived from a choroid plexus papilloma of a human female, is a good epithelial cell model that mimics the *in vitro* characteristics of BCSFB ^11,23,24^. HIBCPP cells retain key epithelial characteristics, including membrane polarity, TJ assembly, energy-sensing kinases, and the expression of transporters and water channel proteins to maintain CSF production and homeostasis in response to metabolic and mechanical stimuli. Studies have demonstrated the expression and polarized localization of TJ proteins, including ZO proteins, claudins, and occludins, as well as ion transporters such as TRPV4, the Na⁺/K⁺-ATPase α1 and β2 subunits, the Na⁺-K⁺-Cl⁻ cotransporter 1 (NKCC1), the sodium–bicarbonate cotransporter (NBCe2), and volume-regulated anion channels (VRAC) at the apical membrane, and the Na⁺/HCO₃⁻ cotransporter (NBCe1) and anion exchanger 2 (AE2) at the basolateral membrane of HIBCPP cells ^11^. Moreover, an electrophysiological study on HIBCPP cells by Hulme et al reported a rapid multiphasic increase in short circuit current (indicative of the net electrogenic ion movement across the cell) and conductance (a measure of transepithelial permeability) upon TRPV4 activation with the agonist (GSK1016790A). This study also demonstrated enhanced apical fluid secretion upon TRPV4 channel activation by GSK1016790A. However, the functional alterations induced after TRPV4 activation did not compromise the spatial organization of junctional proteins such as ZO-1, suggesting that the observed increase in permeability occurs independently of junctional complex disassembly ^11^. Despite this evidence, the precise molecular pathways by which TRPV4 activity modulates TJ organization and epithelial permeability in the choroid plexus remain unclear.

TRPV4-mediated intracellular calcium signaling can activate variety of kinase pathways effectors (phospholipase C, protein kinase C, phosphatidylinositol 3-kinase (PI3K)), suggesting that channel-mediated modulation may occur through phosphorylation-dependent mechanisms ^11,25,26^. One such kinase that our phosphoproteomics study discovered was AMP-activated protein kinase (AMPK) which is a heterotrimeric serine/threonine kinase that functions as a cellular energy sensor and metabolism controller that integrates metabolic and mechanical signals to regulate epithelial polarity, cytoskeletal dynamics, and junctional stability ^27^. AMPK activity is regulated by site-specific phosphorylation events ^28,29^. Phosphorylation of threonine 172 (Thr172) in the activation loop of the catalytic α1-subunit is essential for AMPK activation and is typically mediated by upstream kinases such as liver kinase B1 (LKB1) or calcium/calmodulin-dependent protein kinase kinase β (CaMKKβ) ^28,29^ (**Fig S1**). In contrast, phosphorylation at serine 496 (Ser496) directly by protein kinase A (PKA) and also through feedback inhibition of activated AMPK have been reported to inhibit AMPK activity by preventing Thr172 phosphorylation, and thereby serving as a negative regulatory mechanism ^30,31^ (**Fig S1**). While phosphorylation at Thr172 has been extensively studied, far less is known about Ser496 inhibitory phosphorylation and its effects on AMPK activity, with only a limited number of studies available ^30,31^. In epithelial cells, active AMPK (phosphorylation at Thr172) has been shown to stabilize TJs and adherens junctions under stress conditions, whereas inhibitory phosphorylation at sites like Ser496 (inactive AMPK) can promote junctional disassembly and increase paracellular permeability ^32^. These dual regulatory mechanisms illustrate that modulation of AMPK phosphorylation could serve as a prime link between mechanosensitive signals, such as those initiated by TRPV4 activation, and dynamic control of epithelial barrier function. While AMPK has been implicated in TJ regulation in other epithelial systems ^33^, its potential role as a downstream effector of TRPV4 signaling in the choroid plexus remains unexplored.

In this study, we investigated phosphorylation-dependent signaling pathways downstream of TRPV4 activation in HIBCPP cells ^24,34^, with a particular emphasis on mechanisms governing TJ organization and epithelial permeability. HIBCPP cells were treated with a TRPV4 agonist (GSK1016790A) or antagonist (RN1734) followed by phosphoproteomic analyses to identify downstream signaling and metabolic pathways altered upon TRPV4 modulation. Notably, our study revealed significant upregulation of phosphorylated kinases and TJ proteins involved in maintaining epithelial barrier integrity. Specifically, treatment with a TRPV4 antagonist followed by agonist exposure induced notable changes in ZO-1, whereas agonist treatment alone altered ZO-1, AMPK, and Claudin-7.

TRPV4 activation increased inhibitory phosphorylation of AMPK at Ser496, demonstrating that TRPV4 modulates epithelial barrier function through AMPK-dependent mechanisms. Our findings underscore the importance of AMPK-regulated pathways in epithelial barrier regulation. Elucidating these mechanisms will advance our understanding of how CPe function contributes to CSF homeostasis and how its dysregulation may underline the pathophysiology of hydrocephalus.

## Results

### TRPV4-dependent changes modulate phosphoproteome network in HIBCPP cells

To investigate whether alterations to TRPV4 activity trigger rapid downstream phosphorylation events, we utilized a well-characterized *in vitro* model of human choroid plexus epithelial cells, HIBCPP. Global proteomics and phosphoproteomics mass spectrometry analyses were performed following short-term (10 min) pharmacological manipulation of TRPV4 with TRPV4 agonist (GSK1016790A) and TRPV4 antagonist (RN1734). The 10 minute time frame was chosen to correlate with observed TRPV4 stimulated functional effects on transepithelial electrolyte and fluid movement ^11^. Three experimental conditions were examined in this study: (1) TRPV4 activation (15 nM GSK1016790A vs DMSO diluent), (2) TRPV4 inhibition followed by activation (25 µM RN1734 pre-treatment +15 nM GSK1016790A vs DMSO), and (3) TRPV4 inhibition (25 µM RN1734 vs DMSO) (**Fig 1**). As expected, global proteomics analysis revealed no significant changes to the overall protein abundance across any treatment groups within the defined log_2_ fold change and −log_10_ p-value thresholds, indicating that acute TRPV4 modulation does not alter global protein expression levels (**Fig 2, top panel**). In contrast, phosphoproteomics analysis identified treatment-specific changes in protein phosphorylation, highlighting acute phosphorylation events as a significant downstream response that alter TRPV4 activity (**Fig 2, bottom panel**).

**Fig 1.**
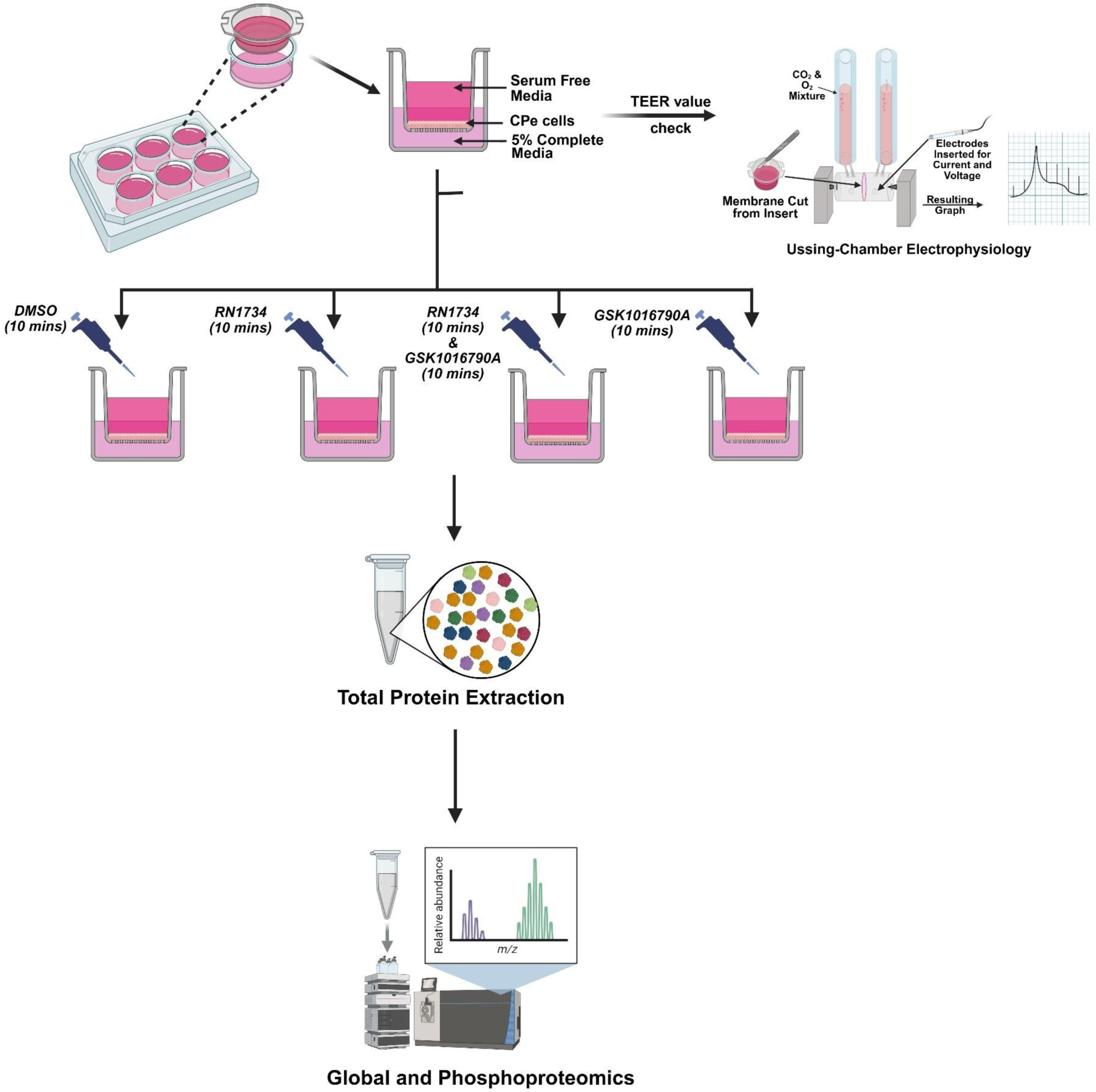
Schematic of experimental setup and drug treatment conditions in HIBCPP cells: Cells were grown on 6-well Transwell inserts with DMEM serum-free media on the apical side and DMEM complete media on the basal side. Transepithelial electrical resistance (TEER) was measured prior to downstream experiments to confirm barrier integrity. For mass spectrometry analysis, cells were treated under four conditions: DMSO for 10 mins, GSK1016790A for 10 mins, RN1734 for 10 mins followed by GSK1016790A for 10 mins, or RN1734 alone for 10 mins, followed by total protein extraction for mass spectrometry. Figure was created using BioRender.

**Fig 2.**
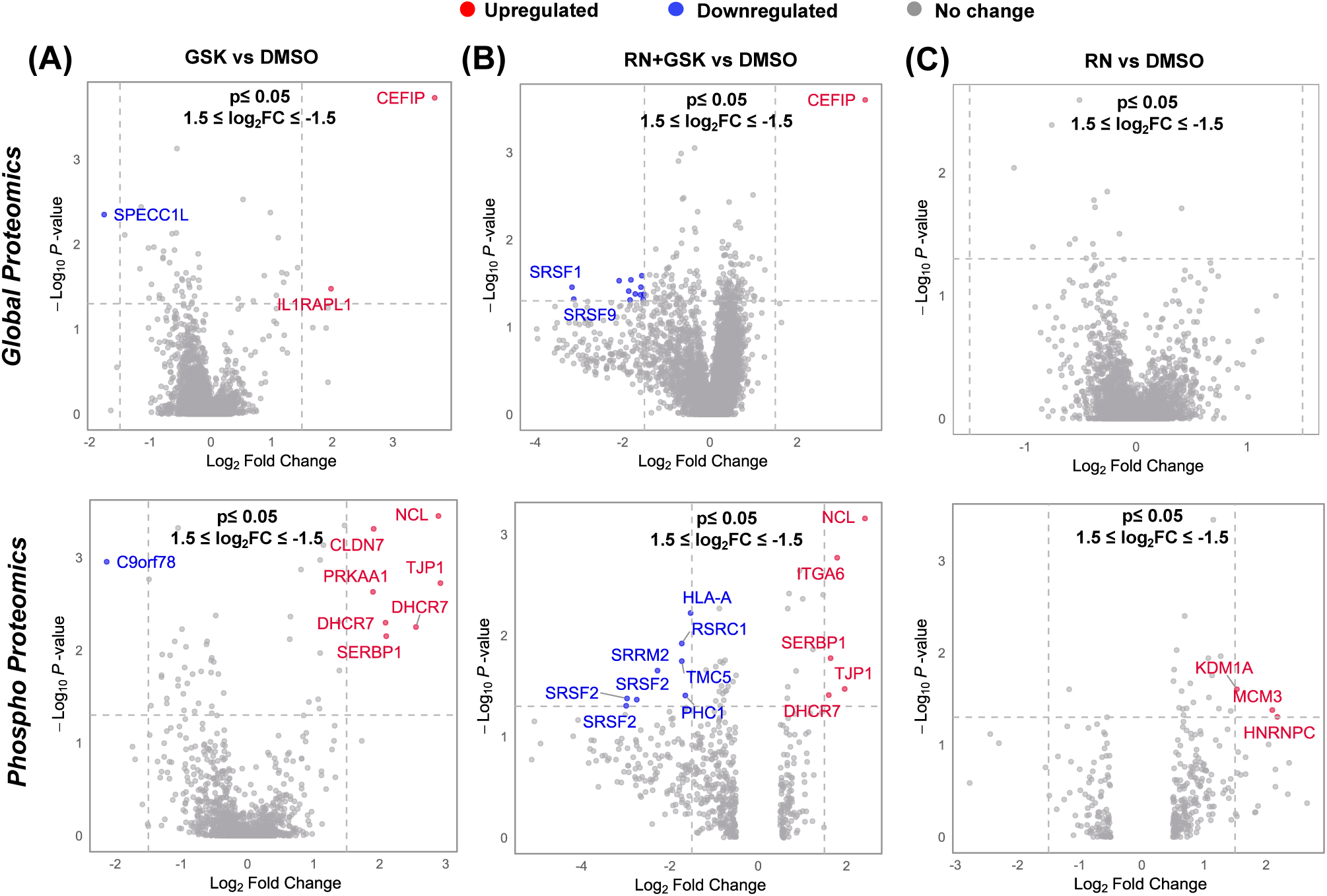
Differentially expressed proteins identified by global (top) and phosphoproteomic (bottom) mass spectrometry analyses: Volcano plots illustrate proteins and phosphoproteins within the thresholds of p < 0.05 and −1.5 ≥ log₂ fold change ≥ 1.5 across three experimental conditions compared with control: (A) TRPV4 agonist (GSK1016790A) vs DMSO, (B) TRPV4 antagonist (RN1734) + TRPV4 agonist (GSK1016790A) vs DMSO, and (C) TRPV4 antagonist (RN1734) vs DMSO. Upregulated, downregulated, and non-significant proteins are shown in red, blue, and grey, respectively. Mass spectrometry analyses were performed using biological triplicates for all conditions (n = 3).

In the first experimental group, in which TRPV4 was activated (GSK1016790A vs DMSO), seven phosphoproteins were differentially regulated, with six showing increased phosphorylation (NCL, CLDN7, PRKAA1, TJP1, DHCR7, and SERBP1) and one showing decreased phosphorylation (C9orf78) (**Fig 2A**). Particularly, these proteins are associated with diverse functions including TJ organization in epithelial cells (CLDN7 or claudin-7 ^35^, TJP1 or ZO-1 ^36,37^), cellular energy homeostasis (PRKAA1 or AMPK ^38^), and RNA binding proteins (NCL ^39^, SERBP1 ^40^), highlighting that TRPV4 activation rapidly influences epithelial barrier-associated signaling and metabolic regulation. In the second experimental group where the cells are first treated with a TRPV4 antagonist and then the agonist (RN1734 + GSK1016790A vs DMSO), eleven phosphoproteins were significantly altered, with five upregulated (NCL, ITGA6, SERBP1, TJP1, DHCR7) and six downregulated (HLA-A, RSRC1, TMC5, PHC1, SRRM2, and SRSF2) (**Fig 2B**). In the third experimental group, the TRPV4 antagonist alone (RN1734 vs DMSO), three phosphoproteins (KDM1A, MCM3, and HNRNPC) exhibited increased phosphorylation (**Fig 2C**).

Among the shared phosphoproteins identified in both the GSK1016790A vs DMSO and RN1734 + GSK1016790A vs DMSO groups, TJP1 (ZO-1), CLDN7 (claudin-7), and PRKAA1 (AMPK) appeared as the top functionally relevant targets based on their established roles in epithelial barrier regulation, membrane organization, and cellular energy sensing (**Fig 3A).** The table in **Fig 3B** summarizes the key functions of the top three candidate phosphoproteins, their corresponding log_2_ fold changes, and a list of other phosphoproteins identified in this study. Further, the heatmap visually represents the phosphorylation patterns of the key phosphoproteins identified across the three experimental groups. Hierarchical clustering highlights distinct phosphorylation signatures corresponding to each experimental condition (**Fig 3C**). The clustering pattern underscores the differential impact of TRPV4 modulation on phosphorylation networks, supporting the concept that TRPV4 activity coordinates a multifaceted phosphoproteomic response in CPe cells.

**Fig 3.**
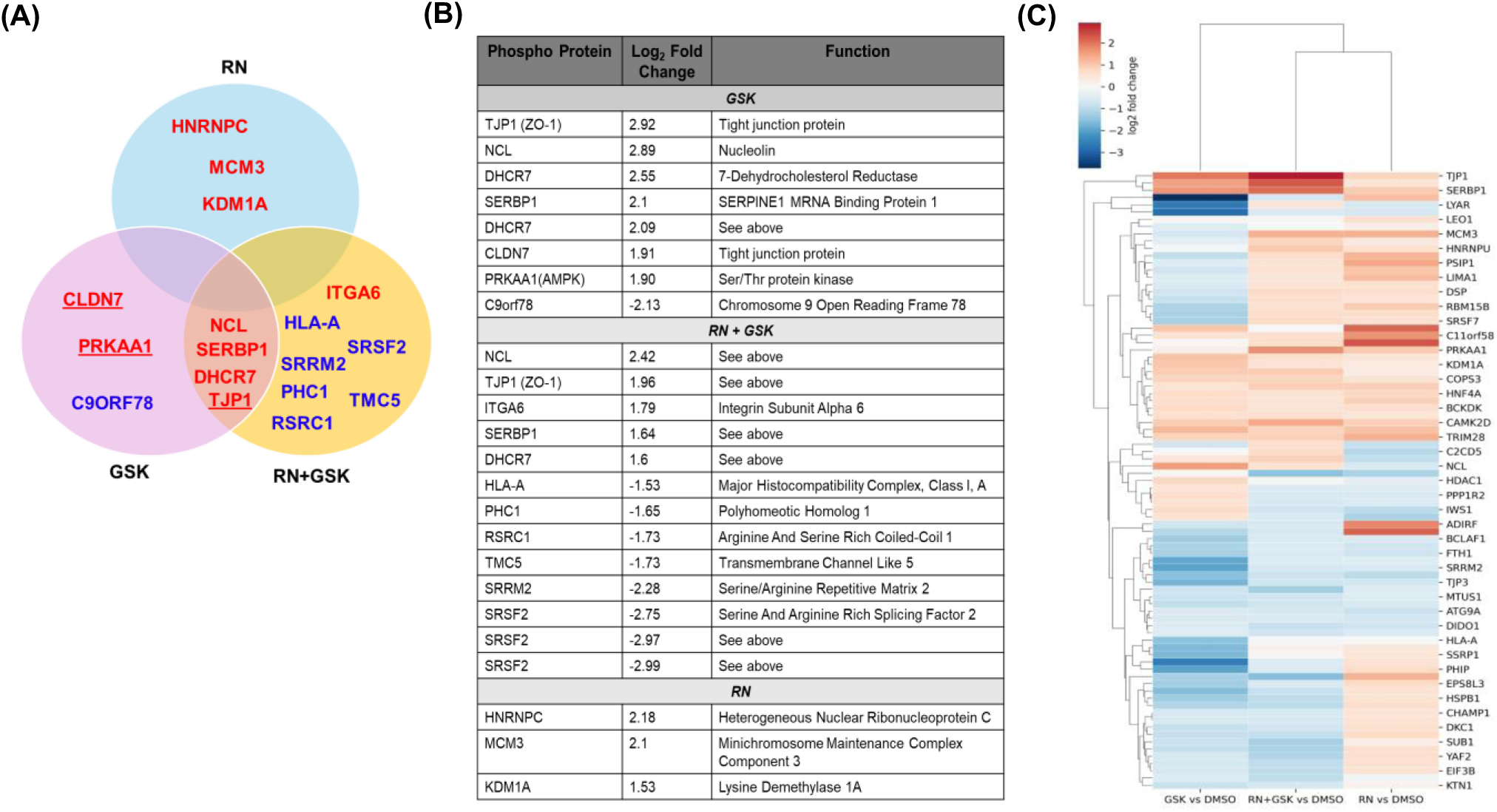
Distinct and shared phosphoproteins differentially expressed across TRPV4 modulation conditions: (A) Venn diagram showing the overlap of significantly altered proteins among GSK1016790A, RN1734, and combined treatment conditions (RN1734+ GSK1016790A) relative to DMSO control. Proteins unique to each condition and those commonly regulated across multiple conditions are indicated, with upregulated proteins shown in red and downregulated proteins shown in blue. Phosphoproteins selected for further study are underlined (B) Table summarizing selected significantly regulated proteins for each comparison, including log₂ fold change values and their annotated biological functions (C) Hierarchical clustering heatmap of differentially expressed proteins across all experimental conditions. Protein abundance values are Z-score normalized and color-coded, with dark red indicating higher expression and blue indicating lower expression relative to DMSO control. Columns represent different experimental conditions (GSK1016790A vs DMSO, RN1734 + GSK1016790A vs DMSO, and RN1734 vs DMSO), and rows represent individual proteins. Heatmap intensities represent averaged measurements from biological triplicates (n=3).

ZO-1 predominantly located in the cytoplasm is a critical scaffolding protein. Phosphorylation of ZO-1 promotes its recruitment from the cytoplasm to TJs, enhancing subcellular localization at the junctional membrane ^41^. This redistribution strengthens TJ assembly and barrier integrity, which is critical for maintaining CPe function ^42^. Calcium-dependent kinases, including AMP-activated protein kinase (AMPK), are known to regulate ZO-1 phosphorylation indirectly through intermediate proteins such as afadin and cingulin, thereby linking intracellular calcium flux to TJ remodeling ^43–45^. Given that RN1734 inhibits TRPV4-mediated Ca²⁺ influx, the persistence of ZO-1 phosphorylation in RN1734 + GSK1016790A vs DMSO suggests that TRPV4 may regulate TJ dynamics in CPe cells through mechanisms that are not solely dependent on Ca²⁺ signaling. Additionally, CLDN7, a transmembrane TJ protein belonging to the claudin family, plays an essential role in maintaining epithelial barrier selectivity and junctional stability ^46^. Phosphorylation of claudins (claudin-1, −3, −4, −5, −7, and −16) has been reported to influence their trafficking, membrane localization, and incorporation into TJ strands. Unlike classical barrier-forming claudins, CLDN7 has a role in maintaining epithelial architecture, cell–matrix interactions, and overall tissue integrity ^47^. It has also been shown to interact with integrins and other membrane proteins, linking TJ organization to cell adhesion and signaling pathways ^48^. As a result, its function extends beyond barrier formation to include regulation of epithelial homeostasis and structural stability.

Next, another crucial phosphoprotein detected in GSK1016790A vs DMSO experimental condition, PRKAA1, the catalytic α1 subunit of AMPK, represents a key metabolic sensor that integrates cellular energy status with epithelial barrier regulation. AMPK activation has been shown to promote TJ assembly and barrier integrity through phosphorylation of junctional proteins, including ZO-1, and through regulation of actin cytoskeletal dynamics ^43^. The phosphorylation of AMPK following TRPV4 activation underscores a functional coupling between TRPV4-mediated calcium signaling and AMPK-dependent metabolic pathways.

To further validate our phosphoproteomic findings, we examined the gene expression of selected TJ proteins including ZO-1, CLDN7, and as well as TRPV4 ion channel in our HIBCPP cells. Reverse transcription PCR (RT-PCR) analysis confirmed the expression of *ZO-1*, *CLDN7*, and *TRPV4* transcripts in HIBCPP cells (**Fig 4A**), verifying that these proteins identified in the phosphoproteomic analysis are transcriptionally expressed in this cell model.

**Fig 4.**
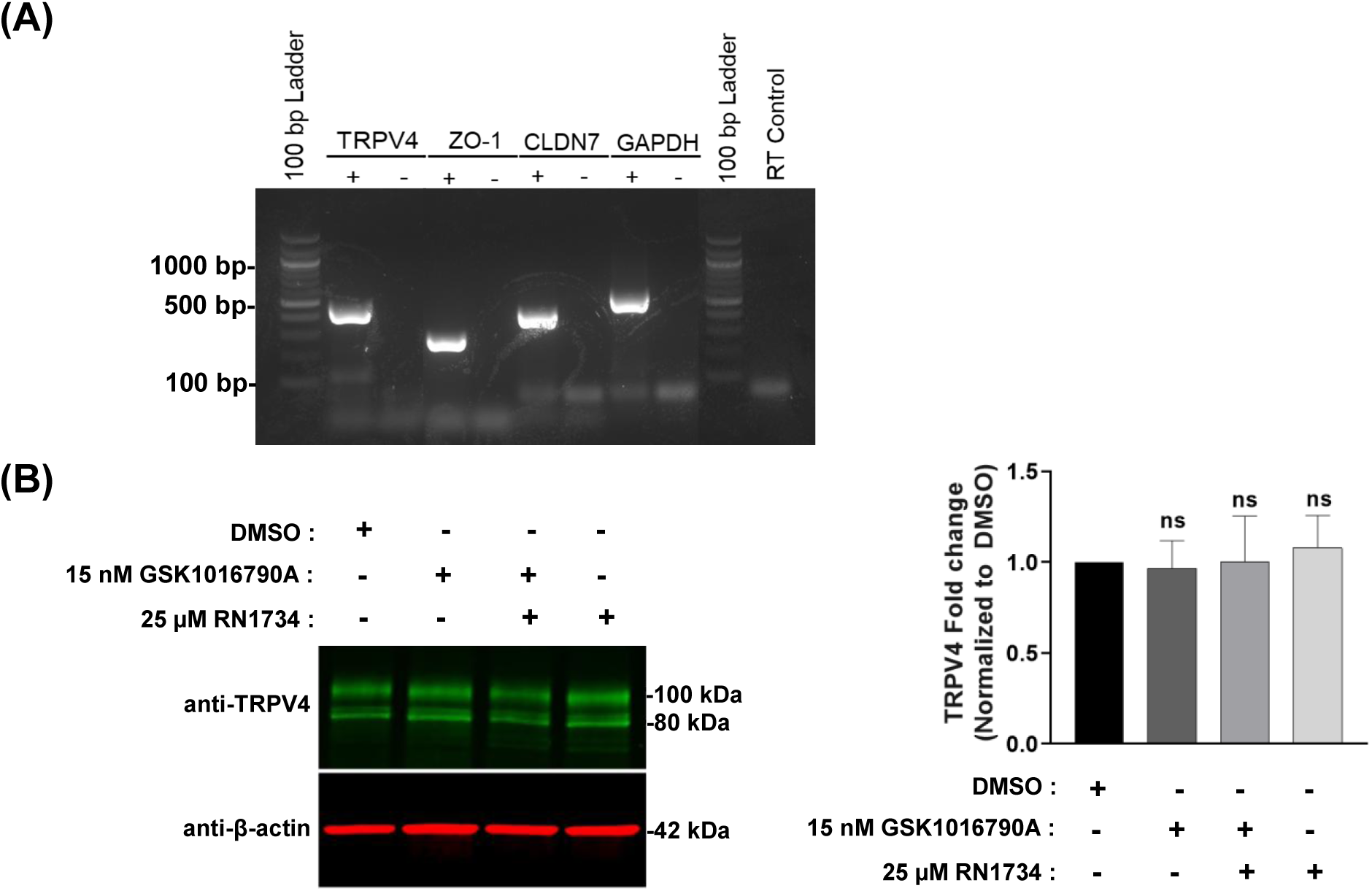
Confirmation of transporter and TJ protein expression in HIBCPP cells: (A) RT-PCR analysis of the transcripts for *TRPV4*, *ZO-1*, and *CLDN7*. RT (reverse transcriptase)-negative controls were included to confirm the absence of genomic DNA contamination, and no-template controls (indicated by “–” under each gene) were included to verify the specificity of amplification. PCR products were visualized on gels stained with SYBR Safe. GAPDH was used as a positive control (B) TRPV4 protein expression was compared across the indicated experimental groups. Western blot analysis revealed two bands at approximately ∼100 kDa and ∼80 kDa. Band intensities were normalized to the housekeeping protein β-actin. Quantification of the bands is shown on the right. Data are presented as mean ± standard deviation from three independent biological replicates (n = 3). Statistical significance was assessed using a two-tailed unpaired Student’s t-test; no significant differences were observed between treatment groups in respect of the DMSO control group (^ns^p > 0.1234).

### TRPV4 protein expression remains constant and localized apically, unaffected by TRPV4 activation and/or inhibition

Global TRPV4 protein levels in CPe cells were analyzed following 10 mins pre-treatment with TRPV4 agonist and antagonist (DMSO for 10 mins, 15 nM GSK1016790A for 10 mins, 25 µM RN1734 for 10 mins + 15 nM GSK1016790A for 10 mins, and 25 µM RN1734 for 10 mins). No significant changes in TRPV4 expression were observed in any experimental group (**Fig 4B**), consistent with the expectation that short-term modulation of TRPV4 activity does not alter overall protein abundance of TRPV4.

Next, we investigated the localization of TRPV4 in CPe cells using immunofluorescence studies to assess its expression and distribution upon activation and inhibition. Previous studies have shown that TRPV4 is predominantly localized to the apical membrane of CPe cells ^11,49^. In our study, we sought to examine whether the localization of TRPV4 is affected by its activation and/or inhibition. Immunofluorescence analysis was performed under four experimental conditions, with TRPV4 expression probed using an anti-TRPV4 antibody. TRPV4 localization remained primarily apical (top quarter of the cells) after treatment with the TRPV4 agonist, antagonist, and a combination of the antagonist followed by the agonist although there were some intracellular expression observed in the bottom quarter closest to the basolateral membrane (**Fig 5**).

**Fig 5.**
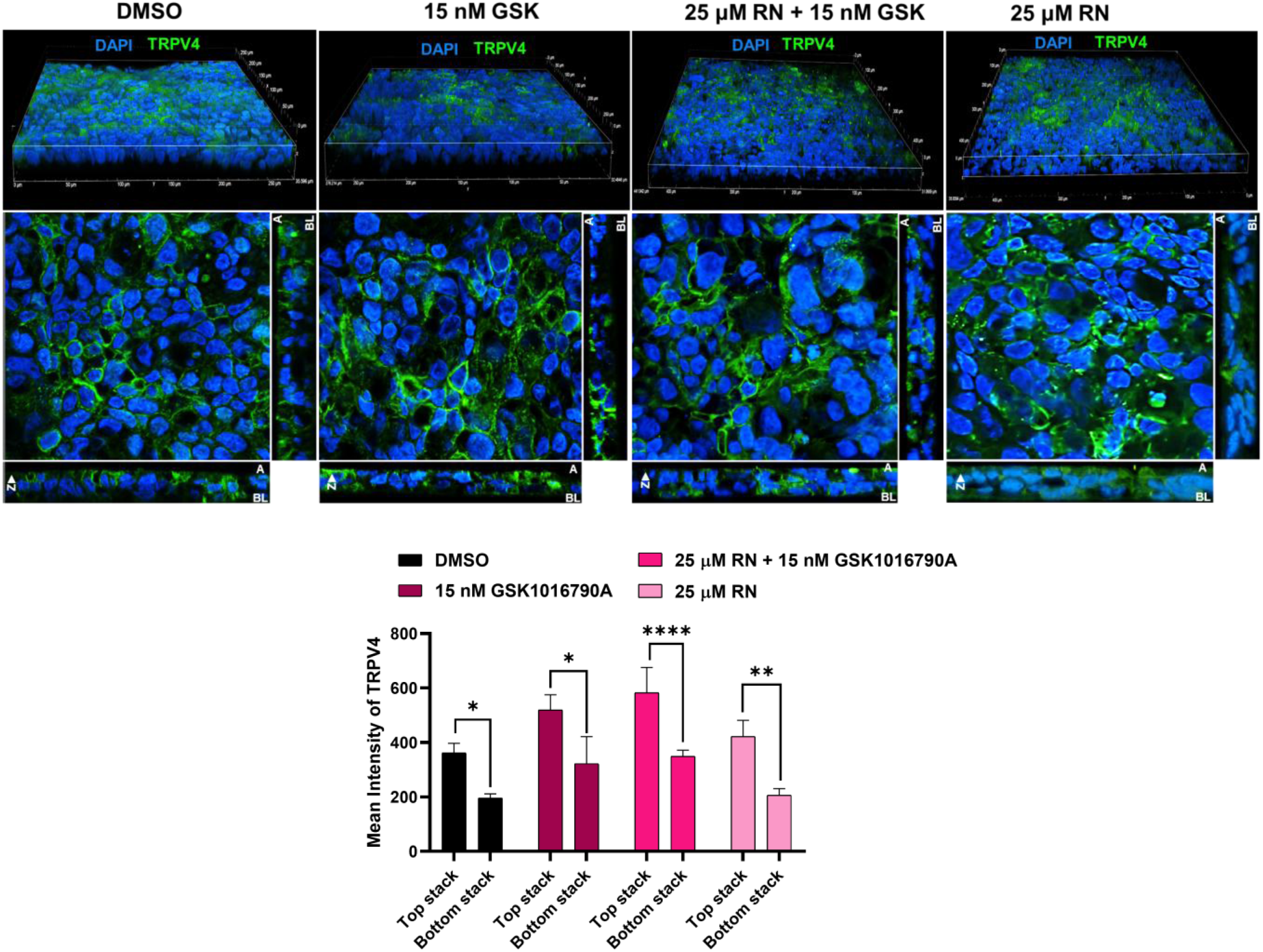
TRPV4 localization in HIBCPP cells: The effect of TRPV4 activation on TRPV4 localization was examined by immunofluorescence. Maximum-intensity projections and three-dimensional (3D) xz-plane renderings of cells grown on permeable supports coated with collagen and stained for TRPV4 (green) and DAPI (blue) are shown under the following conditions: DMSO vehicle control; 15 nM GSK1016790A1016790A for 10 mins; and 25 µM RN1734 for 10 mins followed by 15 nM GSK1016790A1016790A for 10 mins; and 25 µM RN1734 for 10 mins. Z-stack images were quantified as two halves. Top stack represents the top 25 slices, and the bottom stack represents the bottom 25 slices out of 100 slices. Statistical significance was assessed using a two-tailed unpaired Student’s t-test against the GSK1016790Acontrol. Images are representative of three biological replicates (n = 3).

### Pharmacological modulation of AMPK alters TRPV4-mediated barrier permeability

One of the key candidates identified from our phosphoproteomics study was AMPK. As pointed out earlier, the activity of this kinase is primarily controlled by phosphorylation at its two critical sites on the α subunit (Thr172 and Ser496) that act in a coordinated manner. From our phosphoproteomic profiling, AMPK was found to be predominantly phosphorylated at the Ser496 residue in GSK1016790A treated cells (**Table 1**). Additionally, TRPV4 activation also resulted in increased phosphorylation and upregulation of the TJ proteins, ZO-1 and CLDN7, at different serine residues suggesting coordinated regulation of junctional signaling. Given that Ser496 phosphorylation of AMPK may represent a downstream event of TRPV4 channel activation by GSK1016790A, we sought to elucidate the downstream functional changes of this signaling axis.

**Table 1:** Phosphoproteins and corresponding phosphorylation sites identified from our phosphoproteomics study.

| Phosphoprotein | UniProt accession number | Phospho residues |
| --- | --- | --- |
| CLDN7 | O95471 | S204, S206, S207 |
| PRKAA1 | Q13131 | S508, S486, S496 |
| ZO-1 | G3V1L9 | S422, S416, S264, S1710, S427 |

To further investigate the functional consequences of TRPV4-mediated AMPK modulation on epithelial barrier function, we performed Ussing chamber electrophysiological analyses to assess how TRPV4 activation influences AMPK activity, ion transport, and epithelial barrier integrity. AMPK activity was modulated pharmacologically using AMPK activator (AICAr; Millipore Sigma, Cat No: A9978) and AMPK inhibitor (Compound C; Millipore Sigma, Cat No: 171260), followed by the TRPV4 agonist stimulation. Short-circuit current (I_sc_; a measure of net transepithelial electrolyte flux and transepithelial conductance (a measure of permeability or barrier integrity) were monitored under the following conditions: (1) 500 µM AICAr + 15 nM GSK1016790A, (2) 30 µM Compound C + 15 nM GSK1016790A, and (3) 15 nM GSK1016790A alone. Compound C had an inhibitory effect on the basal level of electrolyte flux across the epithelium before the addition of the TRPV4 agonist. Net electrogenic ion transport (I_sc_) stimulated in response to TRPV4 activation was not significantly altered across experimental groups with the exception of a Compound C induced inhibition of transepithelial ion flux in the very early time points (1-2 minutes) of the GSK1016790A response (**Fig 6A**), Conversely, membrane conductance was markedly affected by AMPK modulation. Neither the AMPK activator nor inhibitor had an effect on basal membrane permeability. However, pre-treatment with the AMPK activator, AICAr, appeared to potentiate the TRPV4 mediated electrolyte flux (although it did not show a statistical significance). In contrast, pretreatment with Compound C, the ATP-competitive inhibitor of the AMPK catalytic domain, reduced the TRPV4-stimulated transepithelial conductance significantly, suggesting enhanced barrier integrity/ decreased permeability. (**Fig 6B**).

**Fig 6:**
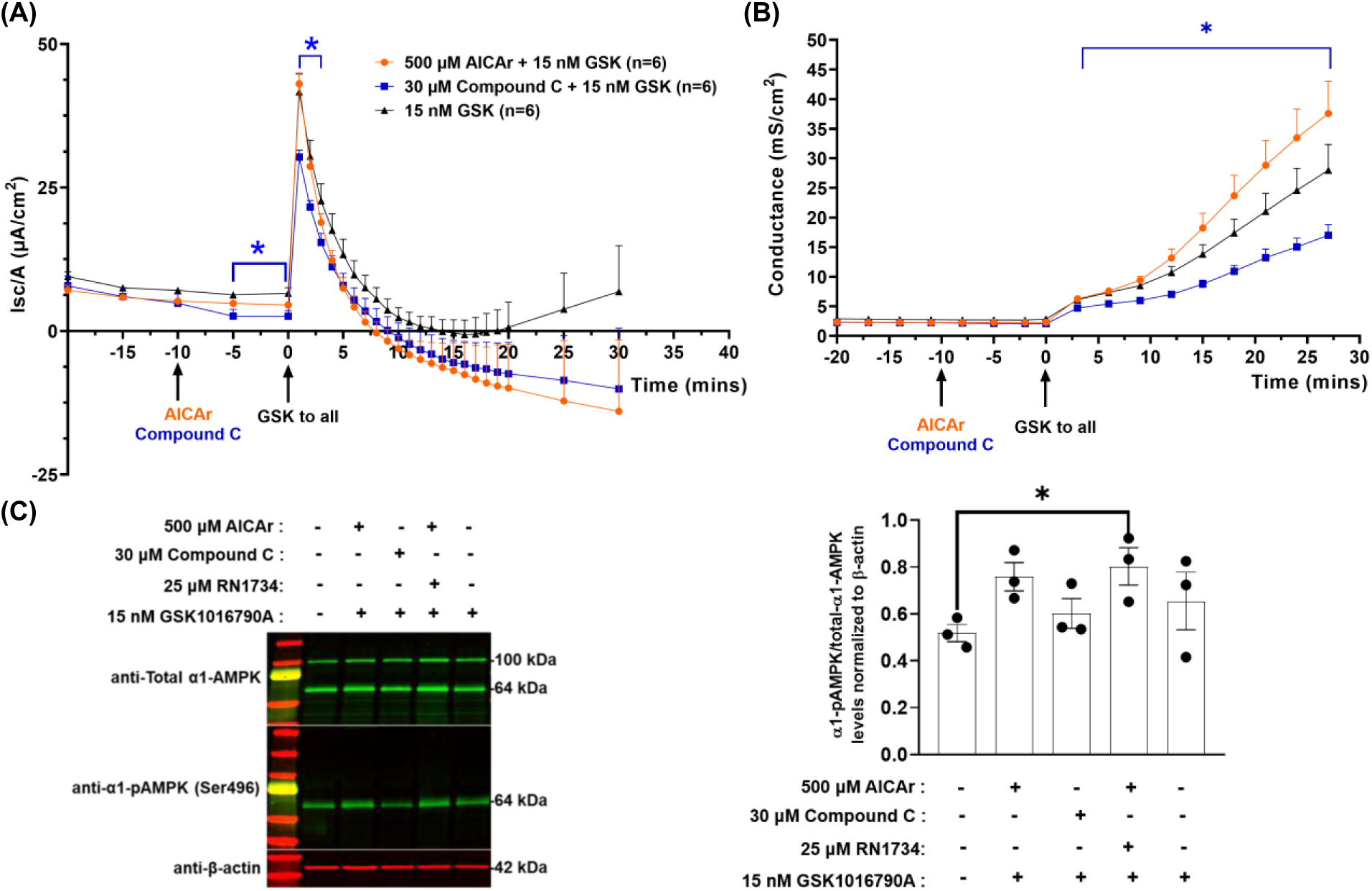
AMPK activation modulates TRPV4-mediated functional effects on cellular permeability: (A) Changes in the short-circuit current (I_sc_) over time were measured by electrophysiology after treatment with AICAr followed by GSK1016790A (dark orange), Compound C followed by GSK1016790A (blue), and GSK1016790A alone (black) (B) Changes in membrane permeability, measured as conductance (mS/cm^2^), were monitored over time after treatment with AICAr followed by GSK1016790A (dark orange), Compound C followed by GSK1016790A (blue), and GSK1016790A alone (black). AICAR, Compound C, and GSK1016790A were added to the apical side only. GSK1016790A was applied to all conditions at t = 0 min, AICAr and Compound C were added at t = −10 mins. The response to GSK1016790A was monitored for 30 mins from t = 0, while responses to the other drugs were monitored for 10 mins. Arrows indicate the time the drugs were applied. Multiple Student’s t-tests were performed in GraphPad Prism 8.0.1. Traces represent standard deviation of six biological replicates (n = 6), and the significance is indicated relative to the GSK1016790A control for each condition (*p < 0.05). (C) HIBCPP cells untreated and treated with AICAr + GSK1016790A, Compound C + GSK1016790A, and GSK1016790A alone were analyzed for total α1-AMPK and α1-pAMPK (Ser496) levels by Western blot (left panel). β-actin was used as the house keeping gene for normalization. Quantification of the blots is shown in the right panel. Individual blot bands for phosphorylated AMPK (α1-pAMPK) to total α1-AMPK levels were normalized to their respective β-actin levels. All values represent the mean ± standard deviation of three biological replicates (n = 3). Statistical significance was assessed using a two-tailed unpaired Student’s t-test. No significant differences were observed in the α1-pAMPK to total α1-AMPK levels amongst the groups except for AICAr + GSK1016790A vs untreated group (*p value < 0.033, ^ns^p > 0.1234).

Furthermore, we validated the protein levels of total α1-AMPK and α1-pAMPK (Ser496) in HIBCPP cells upon AMPK modulation. We utilized the same concentrations and incubation times on cells mentioned in electrophysiological studies and probed with total and phospho (Ser496) AMPK antibodies (**Table S1**). α1-pAMPK (Ser496) to total α1-AMPK ratio varied between treatment groups. Especially, activation of AMPK active form with AICAr followed by GSK1016790A resulted in increased α1-pAMPK to total α1-AMPK ratio when compared to untreated group, suggesting induction of inhibitory AMPK phosphorylation upon AICAr + GSK1016790A treatment. In contrast, inhibition of AMPK inhibitory form with Compound C followed by GSK1016790A reduced α1-pAMPK to total α1-AMPK levels relative to the AICAr group. This finding is consistent with previous studies reporting that Compound C inhibits AMPK kinase activity, thereby reducing Ser496 autophosphorylation^30^. Together with our electrophysiological findings, these results support the notion that reduced inhibitory phosphorylation of AMPK is associated with enhanced barrier function (**Fig 6C**).

These biochemical findings align with our functional data, where elevated and sustained inhibitory phosphorylation of AMPKα1 at Ser496 following AICAr + GSK1016790A treatment likely triggers feedback inhibition of AMPK activity, leading to disrupted epithelial barrier function and increased permeability ^30^. Conversely, inhibition of AMPK inhibitory form with Compound C followed by GSK1016790A reduced Ser496 phosphorylation and corresponded with decreased permeability.

Together, these results support a model in which TRPV4 activation engages AMPK signaling at Ser496 to regulate epithelial barrier properties, such that inhibitory phosphorylation of AMPK compromises barrier integrity, and conversely, reduced inhibitory phosphorylation preserves barrier function.

### Activation of inhibitory phosphorylation on AMPK followed by TRPV4 activation enhances the epithelial barrier discontinuity

In order to further classify the potential roles of AMPK in maintaining barrier permeability, we performed immunofluorescence studies followed by the treatments with AICAr + GSK1016790A, Compound C + GSK1016790A, GSK1016790A alone, AICAr alone, and compound alone and compared it against untreated controls. We monitored the TJ expression of ZO-1 and quantified the junction continuity index (JCI) which measures the degree of fragmentation of the barrier with respect to the drug treatments. Upon treatment with GSK1016790A, the ZO-1 expression appeared discontinuous with a lower JCI than the untreated controls. Interestingly, AICAr treatment followed by GSK1016790A enhanced this more, further decreasing the JCI significantly compared to the untreated controls indicative of more disrupted and less organized ZO-1 expression (**Fig 7**). In contrast, upon Compound C treatment with GSK1016790A, the barrier integrity was preserved almost similar to the untreated controls. Moreover, AICAr treatment alone without GSK1016790A did not cause significant disorganization like what was observed with AICAr with GSK1016790A treatment. Similarly, Compound C treatment alone still maintained the barrier integrity with continuous ZO-1 expression as shown with high JCI values (**Fig 7**). Overall, the ZO-1 expression pattern with treatments indicate that AMPK active form may protect TJ integrity, whereas AMPK inhibitory form may enhance barrier disruption upon TRPV4 activation, consistent with our electrophysiological data.

**Fig 7.**
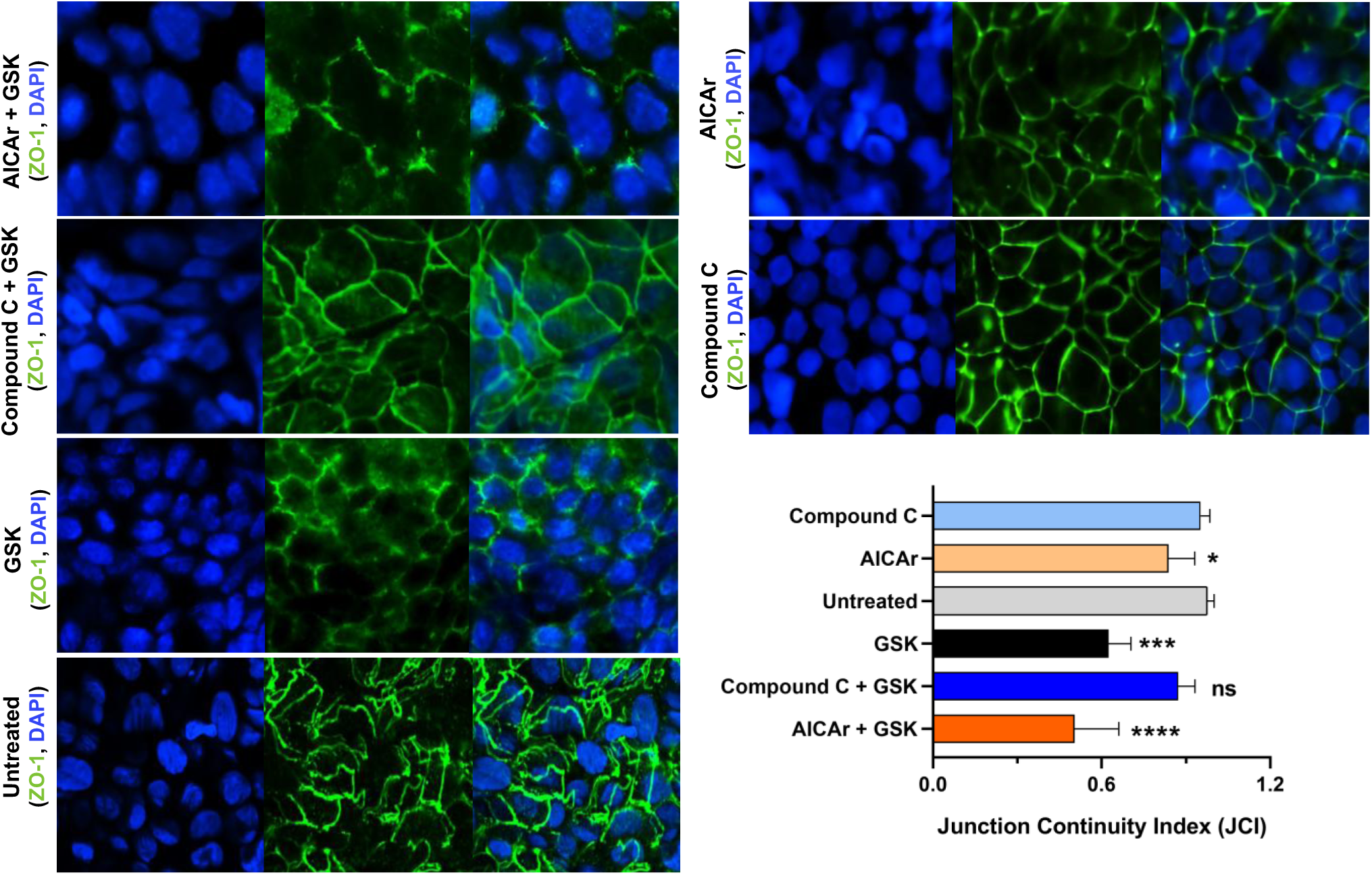
AMPK modulates the TJ integrity: Epithelial membrane integrity was assessed by treating the cells with AMPK and TRPV4 pharmacological modulators (AICAr+GSK1016790A, Compound C+GSK1016790A, GSK1016790A, AICAr, Compound C) and without any treatment (untreated) in immunocytochemistry study. Untreated cells were used as control. ZO-1 (green) expression was quantified to calculate the junction continuity index (JCI). Statistical significance was assessed using a two-tailed unpaired Student’s t-test against the untreated control. Images are representative of three biological replicates (n = 3).

### Metformin-mediated AMPK activation significantly decreased the barrier permeability

Given the regulatory role of AMPK in epithelial barrier function that we are reporting in this study, we used metformin, a well-characterized anti-diabetic drug and an AMPK activator, to evaluate whether modulation of this pathway with metformin influences the observed effects. The cells were treated with 1 mM metformin for 24 hrs at 37 °C followed by TRPV4 activation to measure the transepithelial net electrogenic ion movements and changes to the permeability. Our electrophysiological data showed no significant changes in TRPV4-stimulated transepithelial ion flux (I_sc_) **(Fig. 8A)** but a significant decrease in TRPV4-mediated transepithelial permeabilty upon TRPV4 agonist treatment (**Fig 8B**), implying the metformin-mediated activation of AMPK active form to promote TJ integrity and decreased barrier permeability with TRPV4 stimulation.

**Fig 8.**
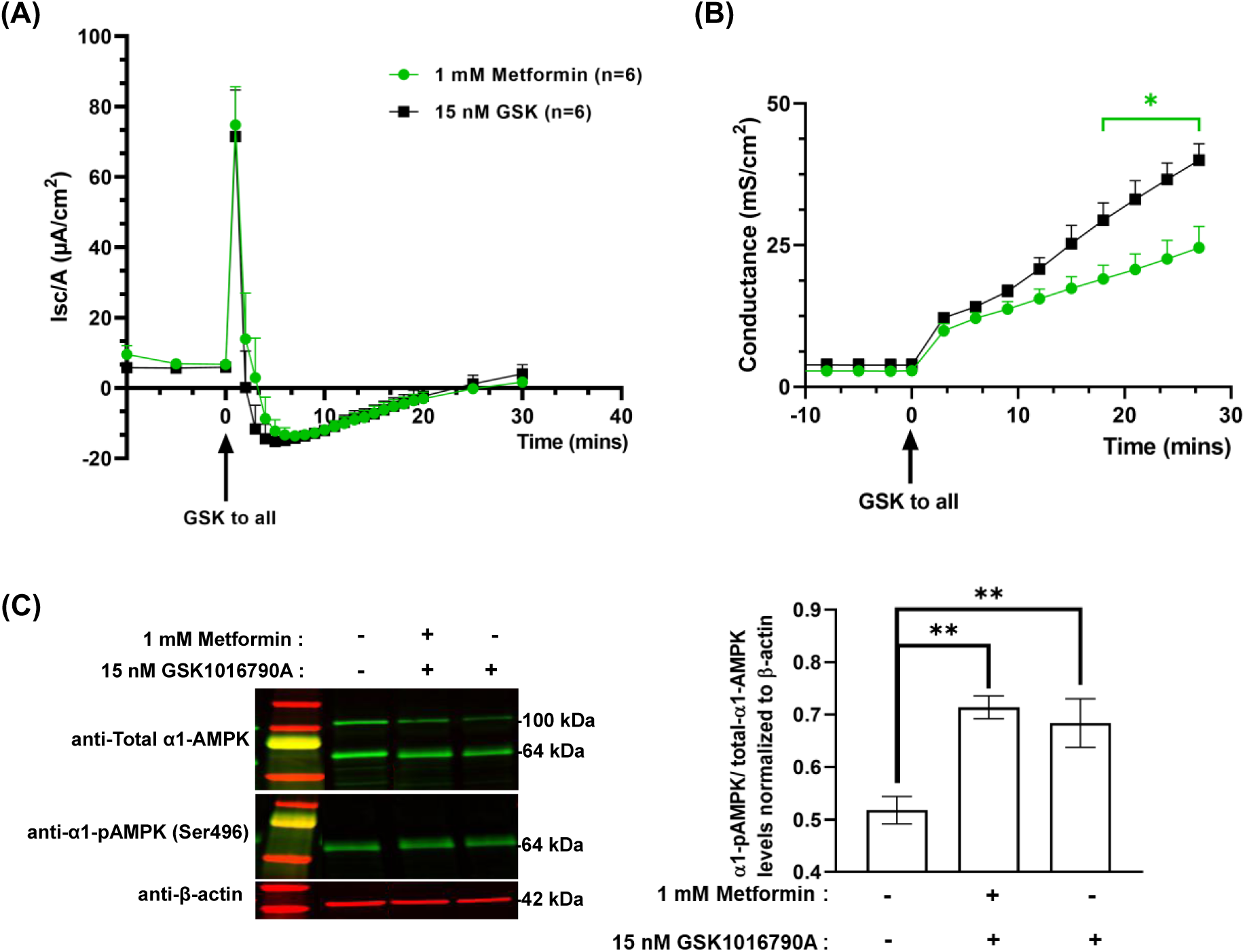
Metformin-Mediated AMPK Activation Regulates Transepithelial permeability: (A) Changes in the short-circuit current (I_sc_) over time and (B) Changes in membrane permeability, measured as conductance (mS/cm^2^), were measured by electrophysiology after treatment with 1 mM Metformin for 24 hours followed by GSK1016790A (green), and GSK1016790A alone (black). Metformin was treated on cells for 24 hrs prior to the experiment. GSK1016790A was added to the apical side of both conditions at t=0. Multiple Student’s t-tests were performed in GraphPad Prism 8.0.1 (*p < 0.05). Traces represent standard deviation (SD) of six biological replicates (n = 6), and the significance is indicated relative to the GSK1016790A control for each condition (C) HIBCPP cells untreated and treated with Metformin + GSK1016790A, and GSK1016790A alone were analyzed for total α1-AMPK and α1-pAMPK (Ser496) levels by Western blot (left panel). β-actin was used as the house keeping gene for normalization. Quantification of the blots is shown in the right panel. Individual blot bands for phosphorylated AMPK (α1-pAMPK) to total α1-AMPK ratio were normalized to their respective β-actin levels. All values represent the mean ± SD of three biological replicates (n = 3). Statistical significance was assessed using a two-tailed unpaired Student’s t-test against the untreated control phospho group (*p value < 0.033, ^ns^p > 0.1234).

Further, total α1-AMPK and α1-pAMPK (Ser496) levels were quantified with metformin + GSK1016790A and GSK1016790A control and compared against untreated controls. Metformin treated cells with GSK1016790A, and GSK1016790A alone group showed higher α1-pAMPK to total α1-AMPK ratio than the untreated controls (**Fig 8C**). As metformin is primarily known to exert its effects through AMPK-dependent signaling pathways, and our electrophysiological data demonstrated reduced epithelial permeability following metformin and GSK1016790A treatment, lower levels of inhibitory AMPK phosphorylation at Ser496 would have been anticipated. Contrary to this expectation, Ser496 phosphorylation was increased. Several mechanisms may explain the observed increase in Ser496 phosphorylation. One possibility is that prolonged metformin exposure resulted in sustained AMPK activation, leading to Ser496 autophosphorylation as a negative-feedback mechanism that limits AMPK activity to some extent. Alternatively, Ser496 phosphorylation may be mediated by another upstream kinase independent of the canonical cAMP/PKA pathway, as Akt has also been reported to phosphorylate this inhibitory residue^50^. These possibilities suggest that multiple regulatory pathways may contribute to the modulation of AMPK activity under our experimental conditions which will need further investigations.

## Discussion

CPe regulates barrier function through highly specialized epithelial transporters, ion channels, and TJ complexes, enabling it to tightly control the composition and movement of CSF. Through coordinated ion transport, water flux, and selective permeability, CPe plays a central role in maintaining CSF homeostasis and intracranial pressure, processes that become dysregulated in conditions such as hydrocephalus. In this context, studies have highlighted the importance of mechanosensitive ion channels in modulating CSF dynamics, particularly through their influence on epithelial transport and barrier properties ^11,51^. Among these, TRPV4 has been implicated as a key regulator of osmosensation and mechanotransduction in CPe, where it contributes to cellular responses to mechanical and osmotic stress. Importantly, a previous *in vivo* study has demonstrated that TRPV4 activation with GSK1016790A can ameliorate ventriculomegaly in hydrocephalic rat models, underscoring its functional relevance in regulating CSF production and ventricular expansion^17^. However, the downstream mechanisms by which TRPV4 mediates these effects remain incompletely understood, particularly in respect to intracellular signaling pathways that link TRPV4-mediated ion flux to epithelial barrier regulation and CSF homeostasis. To address this gap, we investigated TRPV4-dependent signaling using a phosphoproteome-wide approach in human choroid plexus epithelial cell model (HIBCPP cells) with short-term TRPV4 agonist treatment, TRPV4 antagonist pretreatment followed by agonist stimulation, and TRPV4 antagonist alone conditions, enabling systematic assessment of phosphorylation changes associated with both activation and inhibition of TRPV4 signaling. Our analysis identified a subset of differentially phosphorylated proteins that were consistently altered upon TRPV4 modulation, with functional enrichment primarily in AMP kinase-associated signaling networks and TJ-related proteins (ZO-1 and CLDN7) (**Fig 2**), demonstrating that TRPV4 activation triggers coordinated downstream kinase signaling cascades that converge on regulators of epithelial junctional architecture, indicating a potential mechanism by which mechanosensitive signaling may influence TJ integrity and, in turn, contribute to the regulation of CSF dynamics in hydrocephalus.

AMPK is recognized as a central energy sensor and a regulator of epithelial barrier integrity, with multiple studies signifying its role in maintaining TJ stability in both epithelial and endothelial systems ^52–54^. More importantly, AMPK regulates the barrier function through phosphorylation of junctional and cytoskeletal proteins, including components of the apical junctional complexes such as ZO proteins and claudins, thereby promoting TJ assembly and preserving barrier permeability under metabolic or mechanical stress conditions ^55^. Consistent with the established barrier-supportive functions of AMPK, our findings demonstrated an increase in an inhibitory phosphorylation site of AMPK (Ser496) upon GSK1016790A treatment (**Fig 2, GSK1016790A vs DMSO)**, suggesting a potential attenuation of AMPK signaling in response to TRPV4 stimulation, raising the possibility that TRPV4-mediated signaling may counteract AMPK-dependent barrier-supportive pathways in CPe. Our study also demonstrated AMPK modulation with AICAr followed by GSK1016790A treatment which resulted in increased epithelial permeability (although not statistically significant), and higher Ser496 expression while Compound C treatment with GSK1016790A reversed these effects (**Fig 6B**). Our electrophysiological and biochemical data further validated this significant alterations in barrier integrity following AMPK modulation and TRPV4 stimulation.

Notably, the transcellular ion movement did not differ significantly across groups, indicating that the observed changes in conductance are likely driven by paracellular ion flux, as supported by altered TJ integrity reflected in ZO-1 expression. Further, our JCI measurements of ZO-1 expression supported our electrophysiological findings under AMPK and TRPV4 modulation. ZO-1 expression showed increased discontinuity following TRPV4 stimulation with GSK1016790A, which was further enhanced with AICAr + GSK1016790A treatment, indicative of a potential contribution of inhibitory AMPK phosphorylation (Ser496) to compromised barrier integrity. In line with this, increased Ser496 phosphorylation under AICAr + TRPV4-stimulated conditions (**Fig 6C**) may reflect reduced AMPK activity associated with increased barrier leakiness as observed with ZO-1 expression. In contrast, Compound C + TRPV4 stimulation resulted in more continuous ZO-1 expression, with JCI values almost similar to the untreated controls (∼1.0), supporting the role of active AMPK in maintaining barrier integrity and aligning with our electrophysiological observations (**Fig 7**). However, AICAr or Compound C treatment alone did not significantly affect barrier integrity, whereas TRPV4 stimulation with GSK1016790A following AICAr or Compound C disrupted ZO-1 continuity, strongly implicating that AMPK-mediated regulation of TJ integrity is dependent on TRPV4 modulation. This finding would explain the apparent paradox in these studies. Notably AICAr is classically an AMPK activator, and Compound C is an AMPK inhibitor. Therefore, if the mechanism of action was simply AMPK activation, one would expect that AICAr would tighten the junctional complexes and decrease transepithelial conductance while Compound C would do the converse. This is opposite of what we find in our electrophysiology and immunohistochemistry experiments. Clearly TRPV4 activation plays a modulating role, likely by changing the conformation/phosphorylation of AMPK and, thereby, altering the final effect. One possible explanation is that AICAr activates AMPK through its intracellular conversion to the AMP analog (AICAR or ZMP), after its phosphorylation by adenosine kinases and directly binds to AMPK at its γ-subunit. This promotes Thr172 phosphorylation and AMPK activation, while sustained AMPK activation subsequently induce autophosphorylation of the inhibitory Ser496 residue as a negative-feedback mechanism. In contrast, Compound C inhibits AMPK catalytic activity by directly binding to the catalytic site and would therefore be expected to reduce Ser496 autophosphorylation (**Fig 9A**). Thus, the increased Ser496 phosphorylation observed following AICAr treatment and its reduction following Compound C treatment along with TRPV4 agonist are consistent with Ser496 functioning as a feedback regulatory site rather than the initiating mechanism of AMPK activation. Although we feel this is the most likely scenario, other explanations are possible because both AICAr and Compound C have off-target effects.

**Fig 9.**
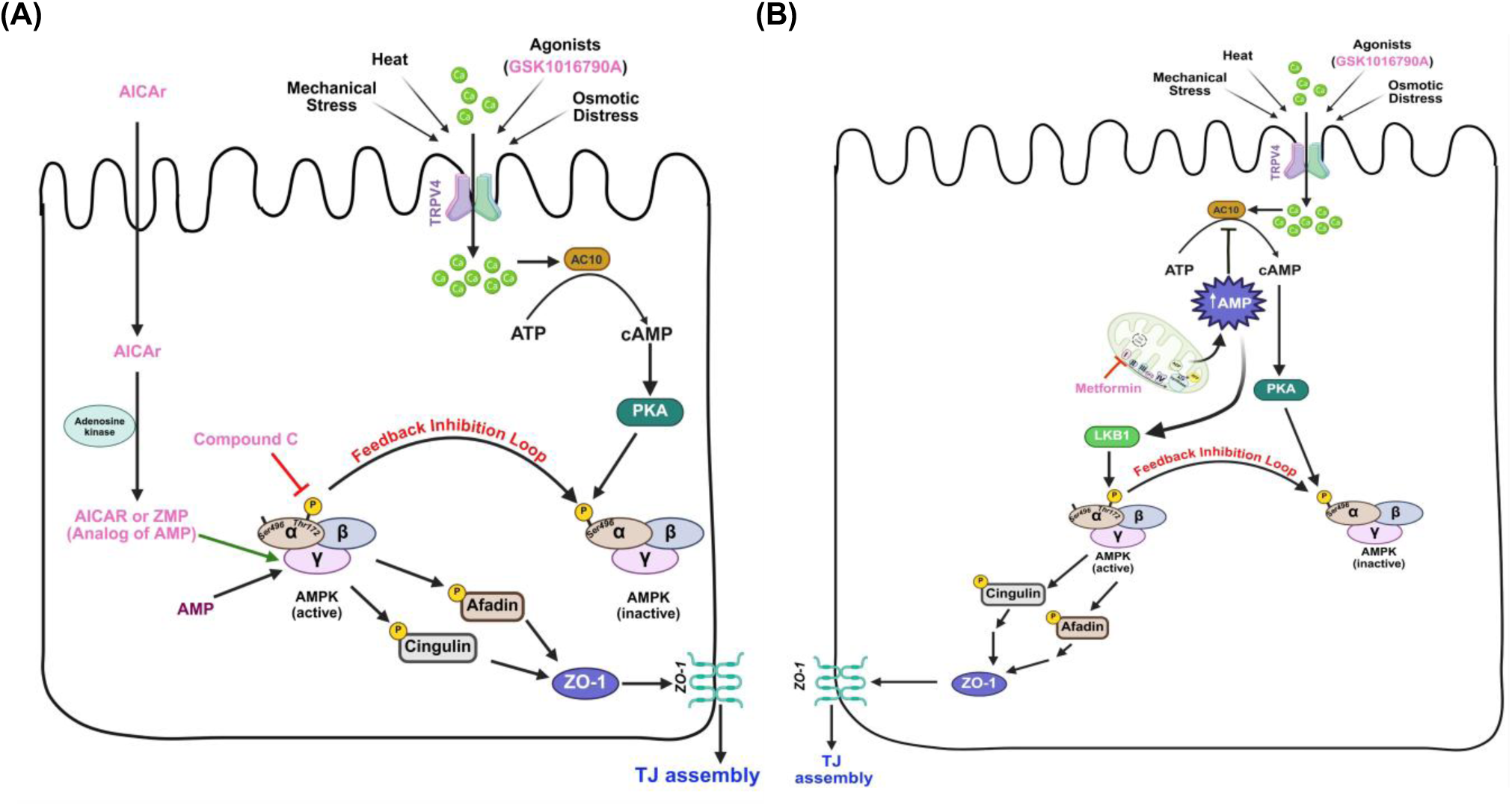
Schematic created in BioRender illustrating the mechanism of action of (A) AICAr, Compound C and (B) Metformin under cellular conditions. Pathways downstream of TRPV4 activation are shown. Protein activation is represented in black arrows, while inhibition is shown in red with blunt-end arrows. Phosphorylation sites reported in the literature and/or observed in our study are indicated (Thr172 and Ser496 in AMPK). Note: AICAr is the cell-permeable nucleoside form of the compound. Following cellular uptake, AICAr is phosphorylated by adenosine kinase to AICAR; also known as ZMP), an AMP analog that binds the AMPK γ subunit and activates AMPK. AC10, adenylate cyclase 10; LKB1, liver kinase b1; PKA, protein kinase A; cAMP, cyclic adenosine monophosphate; ZO-1, zonula occludens-1

Given this apparent interplay between AMPK and TRPV4 signaling, we sought to determine whether pharmacologically available modulators of AMPK would result in similar effects on epithelial barrier function. Emerging literature highlights the role of metformin, a widely used FDA-approved anti-diabetic drug, as an activator of the AMPK pathway ^56^. In epithelial systems, metformin-mediated AMPK activation has been associated with enhanced barrier function, whereas inhibition of AMPK attenuates these protective effects, supporting a functional AMPK dependence ^57,58^. Studies imply that metformin may activate AMPK, further reinforcing its role to probe AMPK-dependent signaling ^59,60^. Metformin exerts its metabolic effects via a different mechanism than AICAr. Metformin acts primarily by inhibiting mitochondrial complex I, leading to an increase in the cellular AMP/ATP ratio and subsequent activation of the LKB1–AMPK signaling axis. LKB1-mediated phosphorylation of AMPK at the Thr172 residue maintains AMPK in its active state, enabling it to regulate cellular energy homeostasis ^59^. Activated AMPK promotes energy-producing (catabolic) pathways while inhibiting energy-consuming (anabolic) processes, thereby enhancing mitochondrial efficiency and overall cellular metabolism. Given this, our study also explored the metformin activity in relation to AMPK-dependent pathway, by performing electrophysiological measurements of net electrogenic ion movement and conductance, to further investigate the AMPK pathway involvement. Although metformin treatment followed by TRPV4 agonist did not alter net transcellular ion movement, conductance was significantly reduced, suggesting enhanced barrier integrity associated with AMPK activation (**Fig 8B**) independent of transepithelial ion flux.

The decrease in conductance observed with metformin treatment is consistent with activation of the LKB1–AMPK pathway in response to increased AMP levels, reflecting a cellular energy stress signal. Under these conditions, AMPK activation may shift signaling away from cAMP/PKA-dependent pathways. Upon TRPV4 activation, which elevates intracellular Ca²⁺, this balance between AMPK and cAMP/PKA signaling may contribute to the observed changes in epithelial barrier function as illustrated by the pathway diagram (**Fig 9B**). As LKB1 phosphorylates AMPK at Thr172 residue, it keeps AMPK in its active form and thereby promotes the TJ integrity and reduced permability ^61^.

Although the activation of LKB1-AMPK pathway can indirectly suppress cAMP/PKA signaling which could prevent the inhibitory phosphorylation on Ser496, our biochemical data shows higher Ser496 in metformin and GSK1016790A treated conditions. This may imply that alternative signaling pathways may be engaged under these conditions in addition to its feedback inhibition loop. One potential explanation is the activation of compensatory kinases, such as Akt (PKB) ^31^, which are also capable of mediating Ser496 phosphorylation. These findings point to a more complex regulatory network and warrant further investigation to understand the signaling mechanisms involved.

Overall, our study provides evidence for the critical role of AMPK-mediated regulation within the TRPV4 signaling axis, indicating that TRPV4-dependent modulation is a key determinant of AMPK activity and downstream control of TJ integrity. Integrating our electrophysiological, biochemical, and imaging data, we propose this mechanistic model (**Fig 9A&B**) in which TRPV4 activation converges on AMPK signaling to regulate epithelial barrier properties, highlighting potential points for pharmacological intervention (given in pink). Thus, targeting TRPV4–AMPK signaling axis may offer a strategy to modulate ion flux across the CPe, thereby influencing CSF dynamics and representing a potential therapeutic avenue in hydrocephalus.

## Methods

### Collagen IV coating of the Transwell insert filters

Cells were grown on Transwell membrane supports that were collagen coated. A concentration of 2 mg/ml stock solution of Collagen IV (Sigma, Cat No: C5533) was prepared by dissolving the powder in ice-cold 0.25% acetic acid. The 2 mg/ml stock solution was diluted in a ratio of 1:40 with sterile MilliQ water in a conical tube to obtain a final working solution of 50 µg/mL. A volume of 300 µL of the working solution was added to the apical side (top) of each insert, followed by gentle shaking or tapping of the plate to ensure that the entire filter was evenly covered by the coating solution.

Transwell plates were placed in the 37 °C, 5% CO_2_ incubator for 1 hour. After 1 hour, the coating solution was carefully aspirated, and the filters were air dried for approximately 20 minutes under sterile conditions. A volume of 3 ml DMEM complete media (composition below) was added on the basolateral side (bottom) of the inserts, and the plates were placed back in the incubator until they were used to seed the epithelial cell cultures.

### Cell culture

HIBCPP cells, a well characterized model of choroid plexus epithelial cells derived from a female cancer patient were used in this study ^24^. Cells were cultured in 25-cm^2^ flasks in Dulbecco Minimal Eagle Medium (DMEM powder, Gibco, Cat No: 12100-046: High Glucose + L-Glutamine + Phenol Red w/ 3.7g/L Bicarbonate) supplemented with 10% fetal bovine serum (Gemini Bio Benchmark, Cat No: 100-106), 1% antibiotic (Penicillin/Streptomycin 100×, Cytiva, Cat No: SV30010), 0.125% insulin (Human recombinant 4 mg/ml in zinc solution, Gibco, Cat No: 12585-014), and 3.7 g/L sodium bicarbonate (Fisher Scientific, Cat No: S233-500) hereafter referred to as “DMEM complete media”. Media in 25-cm^2^ flasks was replenished every other day until the cells reached ∼90% confluency at 37 °C and 5% CO_2_. Cells were detached from flasks using 0.25% sterile-filtered Trypsin-EDTA (Sigma Aldrich, Cat No: T4049) after two washes with Hank’s balanced salt solution (HBSS, Gibco, Cat No: 14170-112) for 5 minutes. Cells were reseeded in 25-cm^2^ flasks to continue until seven passages from the time of thawing and into collagen-coated 6-well Transwell plates for experiments. The Transwell membrane inserts used in these experiments were Costar 0.4 µm polycarbonate membrane tissue culture treated (Corning, Cat No: 3412). Cells seeded onto the Transwell inserts were cultured for 12-16 days, with DMEM complete media for the first 8 days and DMEM serum-free media (DMEM powder, 1% antibiotic (Penicillin/Streptomycin 100×), and 0.125% insulin (Human recombinant 4 mg/ml in zinc solution), and 1.85 g/L sodium bicarbonate) thereafter. Media replenishment for the Transwell until day 8 was done every other day with DMEM complete media and after day 8, cells were fed with DMEM serum-free media every day.

### Protein extraction

HIBCPP cells cultured on the costar Transwell inserts without collagen coating were used in protein extraction studies. Parallel cultures of cells were quickly tested using Ussing-chamber electrophysiology to make sure the TEER value is above 300 Ω·cm^2^ before proceeding with protein extraction. Transwell media were changed to fresh DMEM serum-free media 1 hour before treatment of the cells. After 1 hour, cells were treated with vehicle (DMSO) or relevant drugs (TRPV4 agonist: GSK1016790A1016790A, TRPV4 antagonist: RN1734). Four experimental conditions used in this study for phosphoproteomics mass spectrometry and western blot were (1) DMSO treatment for 10 mins, (2) 15 nM GSK1016790A treatment for 10 mins, (3) 25 µM RN1734 treatment for 10 mins followed by 15 nM GSK1016790A treatment for 10 mins, and (4) 25 µM RN1734 treatment for 10 mins. 3 wells of each condition from a 6-well plate were pooled together to obtain sufficient amount of cells for protein extraction. After treatment, cells were washed with HBSS for 5 mins twice. Cells were then scraped with fresh HBSS and centrifuged at 1200 rpm (145 rcf) for 5 mins. Cell pellets were resuspended in 1× RIPA lysis buffer (Millipore Sigma, Cat No: 20-188) with Halt protease inhibitor cocktail (Thermo Scientific Cat No: 78441). Dissolved pellets were incubated on ice for 30 mins followed by brief sonication for 7 sec, 3 times with 30 sec intervals on ice. After sonication, cells were centrifuged at 12,000 rpm (14,400 rcf) for 20 min at 4 °C and the supernatant was collected and saved as extracted protein from each condition. Protein concentration was quantified using a DC assay kit (Bio-Rad, Cat No: 5000114) with bovine serum albumin (BSA, Bio-Rad, Cat No: 5000206) as standard according to manufacturer’s protocol ^62^. All conditions were performed in triplicate (n = 3).

Western blots were performed using protein extracts obtained from four experimental conditions: (1) untreated cells, (2) treatment with 500 µM AICAR for 10 min followed by 15 nM GSK1016790A for 10 min; (3) treatment with 30 µM Compound C for 10 min followed by 15 nM GSK1016790A for 10 min; (4) treatment with 500 µM AICAR for 10 min followed by 15 nM GSK1016790A for 10 min and subsequently 25 µM RN1734 for 10 min; and (5) treatment with 15 nM GSK1016790A for 10 min alone. All conditions were performed in triplicate (n = 3). Protein concentrations were determined using a DC Protein Assay Kit (Bio-Rad) with BSA as the standard, according to the manufacturer’s instructions.

### Global and phosphoproteomics mass spectrometry

Methods described below are adaptations from literature reports ^63^ and vendor-provided protocols. Samples (n=3 replicates of 4 conditions) were precipitated using trichloroacetic acid (TCA, 20% vol/vol) overnight at 4 °C followed centrifugation at 12,000 rcf for 30 mins and washing of the pellets twice with ice-cold acetone. Proteins were suspended in 8 M urea with 100 mM Tris pH 8.5 and protein concentrations were determined by Bradford protein assay (Bio-Rad, Cat No: 5000006).

Approximately 60 µg equivalent of protein from each sample were then reduced with 5 mM tris(2-carboxyethyl) phosphine hydrochloride (TCEP, Sigma-Aldrich, Cat No: C4706) for 30 mins at room temperature and alkylated with 10 mM chloroacetamide (CAA, Sigma Aldrich, Cat No: C0267) for 30 mins at room temperature in the dark. Samples were diluted with 50 mM Tris HCl, pH 8.5 to a final urea concentration of 2 M for Trypsin/Lys-C based overnight protein digestion at 35 °C (1:50 protease: substrate ratio, Mass Spectrometry grade, Promega Corporation, Cat No: V5072.) Digestions were acidified with trifluoroacetic acid (TFA, 0.5% v/v) and desalted on Sep-Pak® Vac cartridges (Waters^TM^, Cat No: WAT054955) with a wash of 1 mL 0.1% TFA followed by elution in 70% acetonitrile 0.1% formic acid (FA).

#### TMT labeling

All peptide samples (n=3 replicates of 4 conditions) were each labeled with Tandem Mass Tag (TMTpro) reagent per manufactures instructions (0.5 mg per sample ThermoFisher Scientific, TMTpro™ Isobaric Label Reagent Set, Cat No: A44520 Lot ZJ405903) overnight hours at room temperature. Samples were checked to confirm >95% labeling efficiency and then quenched with a final concentration v/v of 0.3% hydroxylamine at room temperature for 15 mins. Labeled peptides were then mixed and dried by speed vacuum. Excess TMTpro reagent was removed from the mix by desalting on a 50 mg Waters Seppak cartridge as described above.

#### Phosphopeptide Enrichment

For phosphoproteomics, the combined, labeled peptide mixture was then applied to a Pierce High-Select™ TiO_2_ Phosphopeptide Enrichment tip (ThermoFisher Scientific, Cat No: A32993). After preparing spin tips as per manufacturer’s instructions, the sample was applied twice, washed, and eluted as per manufacturer’s instructions. The phosphopeptide elution was immediately dried and then resuspended in 25 µL buffer A. For global proteomics, the flowthrough from the phosphopeptide enrichment step was dried down by speed vacuum, reconstituted in 0.1% TFA, and one third was fractionated on Sep-Pak® Vac cartridges using methodology and reagents from Pierce™ High pH reversed-phase peptide fractionation kit (Fractions with 12.5%, 15%, 17.5%, 22%, 22.5%, 25%, 35%, and 70% acetonitrile in 0.1% triethylamine, ThermoFisher Cat No: 84868).

Samples were run (1/8th of each global fraction and 1/5th followed by a tech rep of 1/3rd of each phosphopeptide fraction) on an EASY-nLC 1200 HPLC system (SCR: 014993, ThermoFisher Scientific) coupled to an Eclipse Orbitrap™ mass spectrometer with FAIMSpro interface (ThermoFisher). Each multiplex was run on a 25 cm Aurora Ultimate TS column (Ion Opticks Cat No: AUR3-25075C18) in a 50 °C column oven with a 180-min gradient. The gradient was run from 8-38%B over 160 mins; 30-95% B over 10 mins; held at 80% for 2 mins; and dropping from 95-5% B over the final 5 mins (Mobile phases A: 0.1% formic acid (FA), water; B: 0.1% FA, 80% Acetonitrile (ThermoFisher, Cat No: LS122500)). The mass spectrometer was operated in positive ion mode, default charge state of 2, advanced peak determination on, and Easy IC™ on. Three FAIMS CVs were utilized (−45 CV; −55 CV; −65CV and a technical replicate with −40 CV, −50 CV, and −60 CV) each with a cycle time of 1 sec and identical MS and MS2 parameters. Precursor scans (m/z 400-1600) were done with an orbitrap resolution of 120000, RF lens% 30, 50 ms maximum inject time, standard automatic gain control (AGC) target, minimum MS2 intensity threshold of 2.5e^4^, MIPS mode to peptide, including charges of 2 to 6 for fragmentation with 60 sec dynamic exclusion shared across the cycles, excluding isotopes. MS2 scans were performed with a quadrupole isolation window of 0.7 m/z, 32% HCD collision energy, 50000 resolution, 200% AGC target, dynamic maximum IT, fixed first mass of 100 m/z.

#### Data analysis

Resulting RAW files were analyzed in Proteome Discover™ 2.5.0.400 (ThermoFisher) with a Homo sapiens UniProt reviewed proteome FASTA (downloaded 051322, 20292 sequences) plus common laboratory contaminants (73 sequences). SEQUEST HT searches were conducted with full trypsin digest, 3 maximum number missed cleavages; precursor mass tolerance of 10 ppm; and a fragment mass tolerance of 0.02 Da. Static modifications used for the search were: 1) TMTpro label on peptide N-termini, 2) TMTpro label on lysine (K), and carbamidomethylation on cysteine (C) residues. Dynamic modifications used for the search were 1) oxidation on methionine (M) residues, 2) phosphorylation on serine, threonine or tyrosine residues (S, T, Y), 3) deamidation of asparagine or arginine (Q, N), 4) acetylation on protein N-termini, 5) methionine loss on protein N-termini or 6) acetylation with methionine loss on protein N-termini. A maximum of 3 dynamic modifications were allowed per peptide. Percolator False Discovery Rate was set to a strict setting of 0.01 and a relaxed setting of 0.05. IMP-ptm-RS node was used for all modification site localization scores. Values from both unique and razor peptides were used for quantification. In the consensus workflows, peptides were normalized to total peptide amount.

Quantification methods utilized TMTpro isotopic impurity levels available from ThermoFisher. Reporter ion quantification filters were set to an average S/N threshold of 6 and co-isolation threshold of 30%. Modified peptides were excluded from protein level quantification, normalization and roll-up. Resulting grouped abundance values for each sample type, abundance ratio values; and respective unadjusted p-values (calculated by Protein Abundance ratio with no imputation and ANOVA (individual protein)) from Proteome Discover (ThermoFisher, PD 2.5) were exported to Microsoft Excel.

### Artificial cerebrospinal fluid (aCSF) preparation

Bicarbonate free aCSF was used in the electrophysiological chambers and on cells 1.5 hours prior to experimentation. Fresh aCSF was made fresh daily by mixing solution A (17.3 g/L NaCl, 0.45 g/L KCl, 0.41 g/L CaCl_2_. 2H_2_O, 0.33 g/L MgCl_2_. 6H_2_O) and solution B (0.43 g/L Na_2_HPO_4_.7H_2_O, 0.055 g/L NaH_2_PO_4_. H_2_O) in 1:1 ratio ^64^, in MilliQ water. The solution was pH adjusted to 7.1 using 5M NaOH. After bubbling fresh aCSF with CO₂ for 15–20 minutes, culture medium from the Transwell inserts was removed and replaced with fresh aCSF apically and serum-free DMEM basolaterally. Cells were incubated at 37 °C and 5% CO₂ for 1.5 hours before electrophysiology or subsequent analyses.

### Electrophysiological studies

An 8 chamber electrophysiology instrument (Physiologic Instruments Inc.) was used to measure transepithelial electrical resistance (TEER; the inverse of conductance; a measure of transepithelial permeability) and simultaneous changes in short-circuit current (I_sc_), reflecting net electrogenic ion transport across the epithelium. Cells grown on collagen-coated membrane inserts were used in the electrophysiology experiments. Cells were tested in the electrophysiology system between day 12 to day 16 after seeding onto the membrane inserts, and only those exhibiting TEER values above 300 Ω·cm^2^ were used. Before starting the experiments, cell media was replenished with freshly prepared aCSF on the apical side and DMEM serum-free media on the basolateral side and incubated for 1.5 hours at 37 °C and 5% CO₂. Before measuring cell-containing inserts, background resistance of the system was blanked using inserts with only the membrane (no cells). For measurements, Transwell containing confluent monolayers of cultured cells were placed between the chamber halves (P2315A sliders using chamber slide with the area of 1.26 cm²), and electrodes were placed at their respective ports. The apical and basolateral sides of the chambers were filled with pre-warmed, oxygenated fresh aCSF and the chambers were equilibrated at 37°C, using a temperature-controlled water bath. Chambers were continuously aerated with 95% O₂ and 5% CO₂. Electrodes from the Ussing chamber was connected to the multichannel voltage/current clamp controller and calibration was performed. To compensate for system asymmetries resistance readings from the blank membranes were recorded and later subtracted from the measurements obtained with cell-containing inserts by the system. The tissues were allowed to equilibrate for 10 minutes prior to experimental recording in the system.

Short-circuit current (I_sc_; µA), electrical resistance (TEER; Ohm.cm^2^) and conductance was recorded by the system. The AQUIRE 3.0 data acquisition program (Physiologic Instruments Inc.), connected to the electrophysiology system was used to calculate the TEER (Rt × A), short-circuit current normalized to area (I_sc_/A), and conductance normalized to area (Gt/A), where A represents the surface area of the cell monolayer. Rt was determined from brief current pulses (±10 µA). Resistance values were normalized to the surface area and expressed in Ω·cm². Only TEER values above 300 Ω·cm^2^ was considered for further experimentation. Baseline I_sc_ was monitored continuously, and drugs (AICAR, Compound C, GSK1016790A, and RN1734) were added to the apical side for the required time periods (Refer to the drug treatment groups below). Data were collected and analyzed with AQUIRE 3.0 software.

### Drug treatment groups used in electrophysiological studies

Once the TEER value of the cells were confirmed (> 300 Ω·cm^2^), cultures were allowed to stabilize for 10 mins before adding any drugs in the system. The effect of AMPK on TRPV4 activity was assessed by treating cells on the apical side under the following conditions: 500 µM AMPK activator (AICAR; Millipore Sigma, Cat No: A9978) for 10 mins followed by 15 nM TRPV4 agonist (GSK1016790A) for 30 mins; or 30 µM AMPK inhibitor (Compound C; Millipore Sigma, Cat No: 171260) for 10 mins followed by 15 nM GSK1016790A for 30 mins. To assess the impact of TRPV4 inhibition on AMPK-mediated modulation, cells were treated with 500 µM AICAR for 10 mins, 15 nM GSK1016790A for 30 mins, and 25 µM TRPV4 antagonist (RN1734) for 10 mins. As a control, to evaluate the baseline TRPV4 agonist effects, cells were treated with 15 nM GSK1016790A alone for 30 mins.

For the metformin study, the cells were treated with 1 mM of metformin for 24 hrs in the 37 °C incubator. 15 nM of GSK1016790A was added when the cells were mounted in the electrophysiological system the following day. GSK1016790A alone was added to the control groups. All the electrophysiological experiments were performed with n= 6 unless otherwise mentioned. Statistical significance was evaluated using multiple student’s t-tests in GraphPad Prism 8.0.1.

### Western blot

Total protein extracted from each experimental condition (30 µg) were loaded onto the 4-15% Mini-PROTEAN TGX gels (Bio-Rad, Cat No: 456-8086) with precision plus protein dual color standards ladder (Bio-Rad, Cat No: 1610374) at 120 V for 2 hrs at room temperature. Gels were run in 1× Tris/Glycine/SDS buffer (Bio-Rad, Cat No: 610732). Following electrophoretic separation, proteins were transferred onto nitrocellulose membranes using ice-cold 1× Trans-Blot Turbo transfer packs (Bio-Rad, Cat No: 1704157) in Trans-Blot Turbo transfer apparatus operated with the preconfigured mixed–molecular weight setting (Bio-Rad, Cat No: 1704150). Membrane transfer efficiency and total protein content were checked by staining with Ponceau S (0.1% Ponceau S in 5% glacial acetic acid, ThermoFisher, Cat No: A40000279). Membranes were blocked in 1× Tris-buffered saline (TBS, Bio-Rad, Cat No: 1706435) containing 5% non-fat dry milk, followed by overnight incubation at 4 °C with gentle shaking in the desired primary antibodies diluted in blocking buffer supplemented with 0.01% Tween-20 (1× TBST, Bio-Rad, Cat No: 1610781). The following day, membranes were washed three times for 5 mins in 1× TBST and then incubated with secondary antibodies diluted in 5% milk in 1× TBST for 1 hr (See **Table S1** and **Table S2** for primary and secondary antibody details, respectively). Final washes were performed in 1× TBST, and immunoreactive signals were captured using a Li-Cor Odyssey CLx imaging system. Quantitative analysis of band intensities was done with Image Studio Lite™ image processing software (*LI-COR*, RRID:SCR_013715). For band intensity normalization, β-actin was used as a loading control utilizing the same antibody incubation protocol. All experimental conditions were performed in biological triplicate (n = 3).

### RT-PCR

RNA was isolated from cells cultured as confluent monolayers in 25-cm² flasks using the Monarch Total RNA Miniprep Kit (NEB, Cat No: T2010S). RNA concentration was determined using a Nanodrop 2000 spectrophotometer (ThermoFisher) before cDNA synthesis. cDNA was synthesized from total RNA using LunaScript RT SuperMix Kit (NEB, Cat No: E3010L) and subsequently used for PCR analyses. Primer pairs designed using Primer-BLAST ^65^ and those obtained from the literature ^11^ were used in this RT-PCR study. New primer pairs were ordered from Integrated DNA technologies. Details of the primers used in this study are given in **Table S3**. PCR was performed with an annealing temperature of 62 °C on all primers after optimization. Following PCR, electrophoresis was performed with a 2% agarose gel stained with SYBRSafe Gel Stain (Invitrogen, Cat No: S33102) for *TRPV4*, *ZO-1*, *Claudin-7* and *GAPDH* genes along with no template control and no reverse transcriptase control. Bands were visualized under UV light with a ChemiDoc touch imaging system (Bio-Rad, version 3.0.1).

### Immunohistochemistry

Cells grown on collagen-coated Transwell inserts were used in this study. Cells were used in immunofluorescence studies after confirming their barrier integrity (TEER > 300 Ω·cm^2^). Media were changed to aCSF on top and DMEM serum-free on the bottom 1.5 hours prior to drug treatments. Following the drug treatments, cells were washed with 1× phosphate-buffered saline (PBS) twice and fixed in ice-cold 4% paraformaldehyde (PFA) in 1× PBS for 10 mins. Following fixation, cultures were rinsed three times with 1× PBS (5 mins per wash). The filters were then cut, and sectioned into four pieces, and transferred to 12-well plates for immunostaining. Membranes were permeabilized for 10 mins in buffer containing 0.2% Triton X-100, 3% goat serum, and 1% BSA in 1× PBS, followed by a 30 mins blocking step using 3% goat serum and 1% BSA in 1× PBS. Membranes were subsequently incubated with primary antibodies (anti-TRPV4; Abcam, Cat No: AB314454 at 1:250 dilution, and anti-ZO1; Abcam, Cat No: AB216880 at 1:250 dilution) diluted in blocking buffer overnight at 4°C with gentle agitation. On the following day, membranes were rinsed three times with 1× PBS (5 mins per wash) and then incubated with secondary antibodies (goat anti-rabbit IgG (H+L) Alexa fluor plus 488; Invitrogen, Cat No: A32731TR) prepared in blocking buffer (1:1000 dilution) for 1 hr at room temperature in the dark. Samples were washed thrice with 1× PBS (5 mins each) and subsequently stained with DAPI nuclear dye (0.5 µg/mL in 1× PBS) for 10 mins at room temperature, protected from light. After a final 5 mins wash with 1× PBS, membranes were mounted onto slides using Aqua Poly-Mount (Polysciences Aqua-Polymount, Cat No: 18606-5) and was sealed with coverslips. Imaging was performed using a Nikon AX confocal system to acquire 40X water lens (40X Apo LWD Lambda S) and Z-stack images. No-antibody control was utilized to confirm the non-specific bindings from the secondary antibodies. All experimental conditions were performed in biological triplicate (n = 3). All confocal images were processed in NIS elements AR version 5.30.04.

### Statistical analysis

Quantitative data are shown as the mean ± standard deviation from three independent experiments (n=3), except for electrophysiological studies (n = 6). Statistical significance was assessed using either a two-tailed unpaired Student’s t-test or a two-way ANOVA, performed in GraphPad Prism 8.0.1 (RRID:SCR_002798). Significance thresholds were set at p ≤ 0.05, with the following notation: ****p < 0.0001, ***p < 0.0002, **p < 0.0021, *p < 0.033, and ^ns^p ≥ 0.1234.

## Data availability

The raw and processed mass spectrometry data generated in this study will be deposited in the ProteomeXchange repository. All other data are provided within the manuscript and in supplementary file (.xlsx file)

## Ethics approval and consent to participate

Not applicable

## Consent for publication

Not applicable

## Author contributions

G.M. and B.B.Y.: conceptualization, study design, methodology, methodology, investigation, data curation, writing, and editing. G.M.: sample preparation, mass spectrometry analysis, bioinformatics, and data evaluation in collaboration with the center for proteome analysis at the Indiana University School of Medicine (IUSM). B.B.Y supervision, project administration, and funding acquisition. All authors have read and approved the final manuscript.

## Acknowledgements

The mass spectrometry work performed in this study was carried out by the Indiana University School of Medicine (IUSM), Center for Proteome Analysis Core. The authors would like to thank the Center for Proteome Analysis, especially Dr. Emma Doud and Dr. Amber Mosley, for performing the mass spectrometry analyses and acquiring the data used in this study. Acquisition of the IUSM Proteomics instrumentation used for this project was provided by the Indiana University Precision Health Initiative.

