## Supplementray File for "Phosphoproteomic Insights into TRPV4–AMPK Signaling Axis in the Choroid Plexus Epithelium: Implications for Therapeutic Targeting in Hydrocephalus"

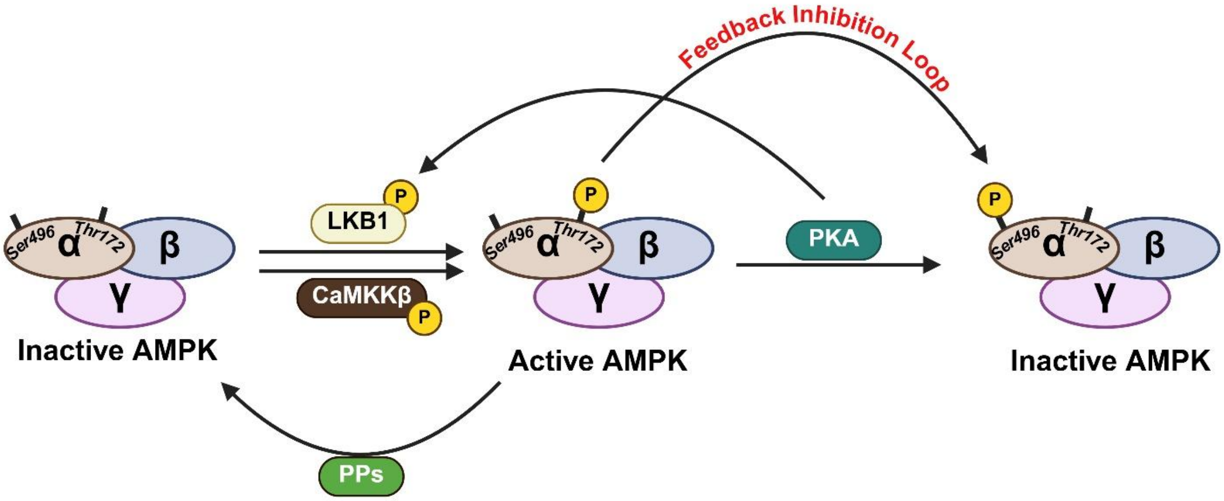


Fig S1 | Dynamic regulation of AMPK activity: AMPK activation is mediated by phosphorylation of Thr172 on the α subunit by upstream kinases LKB1 and CaMKKβ, whereas dephosphorylation by protein phosphatases (PPs) and inhibitory regulation by protein kinase A (PKA) promote AMPK inactivation while activated AMPK also phosphorylates at Ser496 through autophosphorylation to result in feedback inhibition. These opposing mechanisms tightly control AMPK activity in response to cellular signaling and metabolic status.

Table S1: Primary antibodies used in western blot studies

| Primary antibody | Species and clonality | Manufacturer | Antibody dilution |
|---|---|---|---|
| TRPV4 | Rabbit recombinant multiclonal | Abcam (AB314454) | 1:1000 |
| AMPK | Rabbit polyclonal | Abcam (AB3759) | 1:1000 |
| p-AMPK (S496) | Rabbit monoclonal | Abcam (AB92701) | 1:1000 |
| β-actin | Rabbit polyclonal | Proteintech (20536-1-AP) | 1:1000 |

Table S2: Secondary antibodies used in western blot studies

| Secondary antibody | Manufacturer | Antibody dilution |
|---|---|---|

| | | |
|---|---|---|
| Donkey anti-rabbit IgG (H+L) (Alexa Fluor-790) | Jackson Immunoresearch (711-655-152) | 1:4000 |
| Donkey anti-mouse IgG (H+L) (Alexa Fluor-680) | Jackson Immunoresearch (715-625-150) | 1:4000 |

Table S3: Primer sequences used in RT-PCR study

| Gene Name | Forward primer (5'-3') | Reverse primer (5'-3') | Product size (bp) |
|---|---|---|---|
| *TRPV4* | AAGTTCAAGGACTGGGCCTATG | ACGATCACCAGGACAGAGTAGA | 473 |
| *AQP1* | AGATCAGCATCTTCCGTG | AGTTGTGTGTGATCACCG | 354 |
| *TJP1* (ZO-1) | GAGGACCAGCTGAAGGACAG | GGCCACTTCTTGGATCATGT | 243 |
| *CLDN7* (Claudin-7) | CTCGAGCCCTAATGGTGGTC | GCAAGGAGATCCCAGGTCAC | 414 |
| *GAPDH* | AAGCCTCAAGATCATCAGCAA | CCCTGTTGCTGTAGCCAAATTC | 546 |
